# Corticosteroid and progesterone transactivation of mineralocorticoid receptors from Amur sturgeon and tropical gar

**DOI:** 10.1101/053199

**Authors:** Akira Sugimoto, Kaori Oka, Rui Sato, Shinji Adachi, Michael E. Baker, Yoshinao Katsu

## Abstract

The response to a panel of steroids by the mineralocorticoid receptor (MR) from Amur sturgeon and tropical gar, two basal ray-finned fish, expressed in HEK293 cells was investigated. Half-maximal responses (EC50s) for transcriptional activation of sturgeon MR by 11-deoxycorticosterone, corticosterone, 11-deoxycortisol, cortisol and aldosterone, and progesterone were between 13 pM and 150 pM. For gar MR, EC50s were between 8 pM and 55 pM. Such low EC50s support physiological regulation by these steroids of the MR in sturgeon and gar. Companion studies with human MR and zebrafish MR found higher EC50s compared to EC50s for sturgeon and gar MR, with EC50s for zebrafish MR closer to gar and sturgeon MR than was human MR. For zebrafish MR, EC50s were between 75 pM and 740 pM; for human MR, EC50s were between 65 pM and 2 nM. In addition to progesterone, spironolactone and 19nor-progesterone were agonists for all three fish MRs, in contrast to their antagonist activity for human MR, which is hypothesized to involve serine-810 in human MR because all three steroids are agonists for a mutant human Ser810Leu-MR. Paradoxically, sturgeon, gar and zebrafish MRs contain a serine corresponding to serine-810 in human MR. Our data suggests alternative mechanism(s) for progesterone, spironolactone and 19nor-progesterone as MR agonists in these three ray-finned fishes and the need for caution in applying data for progesterone signaling in zebrafish to human physiology.

## INTRODUCTION

The mineralocorticoid receptor (MR) is a transcription factor that belongs to the nuclear receptor family, a diverse group of transcription factors that also includes receptors for androgens (AR), estrogens (ER), glucocorticoids (GR) and progestins (PR), and other small lipophilic ligands, such as thyroid hormone and retinoids [1-5]. The MR and GR are descended from a common corticosteroid receptor (CR), which are present in lampreys and hagfish [5-7]. Several corticosteroids (Figure 1), including aldosterone (Aldo), cortisol (F), 11-deoxycortisol (S), corticosterone (B) and 11-deoxycorticosterone (DOC), as well as progesterone (Prog), are transcriptional activators of Atlantic sea lamprey CR and hagfish CR [6]. Among these steroids, Aldo, the main physiological activator of the MR in human and other terrestrial vertebrates [8-11], had the lowest half-maximal response (EC50) for transcriptional activation of the CR. This strong response to Aldo is surprising because Aldo is not found in either lamprey or hagfish serum [6]. S, which along with DOC is present in Atlantic sea lamprey serum, has been found to have mineralocorticoid activity in lamprey [12, 13].

**Figure 1.**
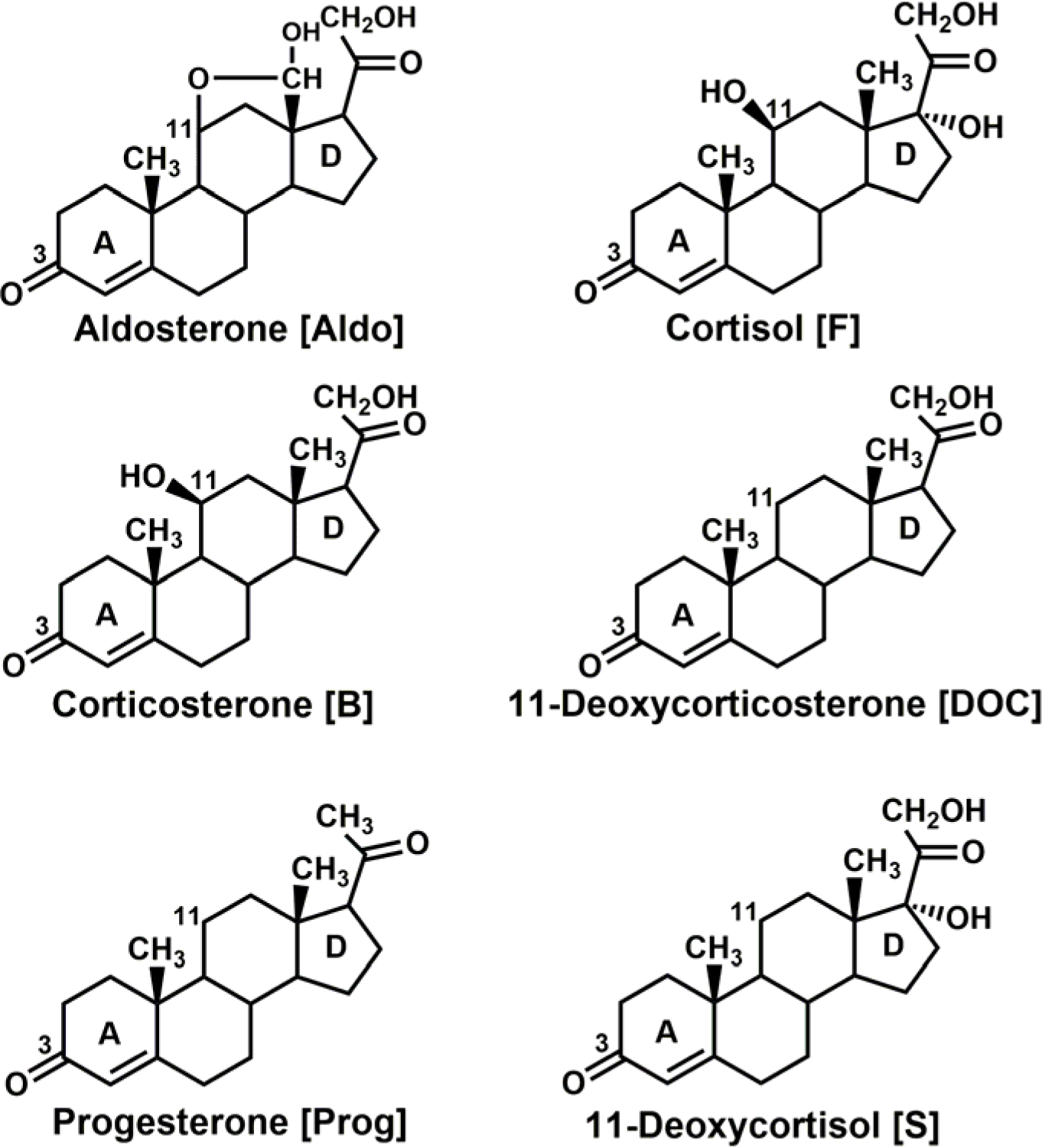
Structures of potential steroid regulators of fish MR. Aldo, the physiological ligand for terrestrial vertebrate MRs, is not found in fish [20]. F and DOC have been proposed to be mineralocorticoids in teleosts [22, 25]. S is a ligand for corticosteroid receptor in lamprey [12]. Progesterone is an antagonist for human MR [27].

Distinct MR and GR genes first appear in cartilaginous fishes (Chondrichthyes), such as sharks, rays and skates [6, 14]. Carroll et al. [14] determined EC50s of several corticosteroids for skate MR; EC50s were 70 pM for Aldo, 30 pM for DOC, 90 pM for B, 1 nM for F and 22 nM for S. In teleosts, which comprise about 95% of known ray-fish species (*Actinopterygii*), corticosteroid activation of the MR has been investigated for cichlid [15], trout [16], carp [17], midshipman fish [18] and zebrafish [19], with Aldo, F and DOC being the principal steroids that were studied. Although Aldo has not been found in teleost fish [20], Aldo has a low EC50 for teleost MRs, similar to that found for Aldo activation of lamprey CR and skate MR. DOC also has a low EC50 for teleost MRs, and DOC has been proposed as mineralocorticoid in fish [16, 21-25]. F also has been proposed to be ligand for teleost fish MR [22, 24, 25]. The response of the teleost MRs to B and S, which are found in fish [25, 26], has been studied only in trout, in which the EC50s are 10 nM for B and 3.7 nM for S [16]. Interestingly, spironolactone (spiron) and Prog are agonists for trout MR [16], in contrast to their antagonist activity for human MR [16, 27]. Spiron also is an agonist for zebrafish MR [19]. Together, these studies indicate that several corticosteroid(s) are potential transcriptional activators of teleost MRs [22, 25, 28, 29].

An important gap in our understanding of the evolution of selectivity of ray-finned fish MRs for steroids is the absence of data on the MR in Chondrostei (sturgeons, paddlefishes, reedfishes, bichirs) and Holostei (bowfins, gars), which evolved before a fish-specific genome duplication occurred after the split of the Acipenseriformes (sturgeons) and the Semionotiformes (gars) from the lineage leading to teleost fish, but before the divergence of Osteoglossomorpha (Figure 2) [30-32].

**Figure 2.**
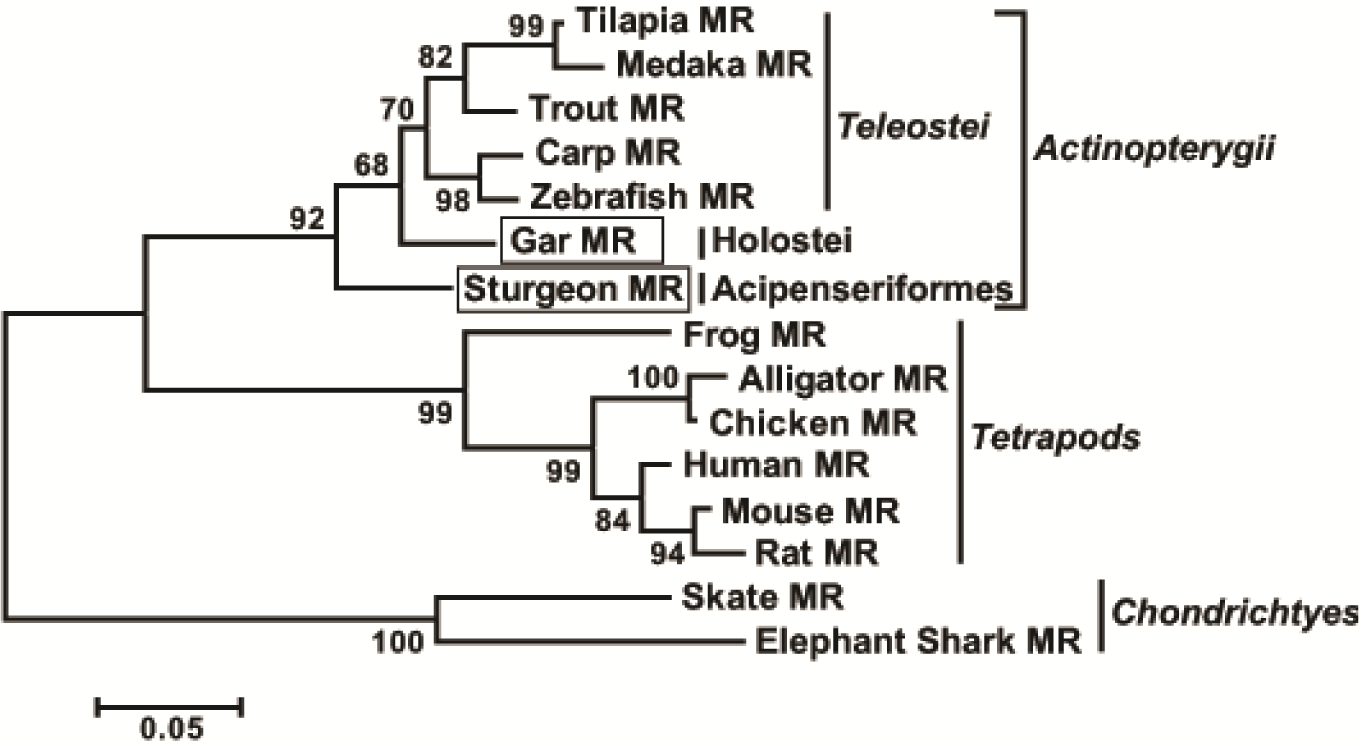
Phylogenetic relationship of sturgeon and gar MRs to other vertebrates. To investigate the relationship of sturgeon and gar to other fish, we constructed a phylogenetic tree of the steroid-binding domains on MRs in sturgeon, gar, selected teleosts, elasmobranchs and tetrapods. The phylogenetic tree was constructed using the maximum likelihood with JTT+G model with 1000 bootstrap replications, which are shown as percentages at the nodes of the tree.

Our interest in the evolution of steroid hormone action [4, 5, 33, 34] prompted us to investigate transcriptional activation of the MR from Amur sturgeon, *Acipenser schrenckii*, and tropical gar, *Atractosteus tropicus* by a broad panel of corticosteroids (Aldo, F, B, DOC, S), Prog, spiron and 19nor-progesterone (19norP), a steroid that has not previously been studied for activation of fish MR. To gain further insight into the evolution of steroid specificity, we compared our results with companion studies of zebrafish and human MRs. In agreement with studies of teleost MRs, we find that Aldo and DOC have the lowest EC50 (highest activity) for sturgeon and gar MRs. However, we also find that S, B, F, and Prog have low EC50s, consistent with these steroids also having a physiological role as ligands for these MRs. Spiron and 19norP also activated sturgeon and gar MRs. In comparison, zebrafish MR has a strong response to Aldo and DOC and a good response to B, F, S Prog, spiron and 19norP, while human MR has strong response to Aldo, DOC and B and a good response to F and S, and a weak response to Prog and 19norP and no response to spiron. The weak response to Prog and 19norP and absence of a response to spiron by human MR is in agreement with other studies [27, 35, 36]. The strong response to 19norP, Prog and spiron of sturgeon, gar and zebrafish MR is perplexing because the basis for the low response to to these steroids by human MR is thought to be due to the presence of Ser-810 on α-helix 5 [27, 37, 38]. 19norP, Prog and spiron are agonists for human MR with Ser810Leu mutation [27, 37, 38]. Sturgeon, gar and zebrafish MRs contain a serine corresponding to serine-810 in human MR, suggesting the presence of an alternative mechanism for these steroids acting as MR agonists in these three ray-finned fishes, as well as the need to apply caution in interpreting data on Prog activity in zebrafish to human physiology.

## MATERIALS AND METHODS

### Animals and chemical reagents

Amur sturgeon and tropical gar were obtained as described previously [32]. All experimental procedures involving live fish followed the policies and guidelines of the Hokkaido University Animal Care and Use Committee. Aldosterone (Aldo) (CAS 52-39-1), corticosterone (B) (CAS 50-22-6), cortisol (F) (CAS 50-23-7), 11-deoxycortisol (S) (CAS 152-58-9), 11-deoxycorticosterone (DOC) (CAS 64-85-7), progesterone (Prog) (CAS 57-83-0), 19nor-progesterone (19norP) (CAS 472-54-8), spironolactone (Spiron) (CAS 52-01-7), 5α-dihydrotestosterone (DHT) (CAS 521-18-6), and 17β-estradiol (E2) (CAS 50-28-2) were purchased from Sigma-Aldrich. For the reporter gene assays, all hormones were dissolved in dimethyl-sulfoxide (DMSO) and the final concentration of DMSO in the culture medium did not exceed 0.1%.

### Molecular cloning of mineralocorticoid receptors

Two conserved amino acid regions, GCHYGV and LYFAPD of vertebrate MRs were selected and degenerate oligonucleotides were used as primers for PCR. First-strand cDNA was synthesized from 2 µg of total RNA isolated from the liver after amplification, and an additional primer set (CKVFFK and LYFAPD) was used for the second PCR. The amplified DNA fragments were subcloned with TA-cloning plasmid pGEM-T Easy vector, sequenced using a BigDye terminator Cycle Sequencing-kit with T7 and SP6 primers, and analyzed on the 3130 Genetic Analyzer (Applied Biosystems). The 5’- and 3’-ends of the mineralocorticoid receptor cDNAs were amplified by rapid amplification of the cDNA end (RACE) using a SMART RACE cDNA Amplification kit.

### Database and sequence analysis

MRs for phylogenetic analysis were collected with Blast searches of GenBank. A phylogenetic tree for MRs was constructed by the Neighbor-Joining Method [39] after sequences were aligned by MUSCLE [40] using several fish, frog, alligator, chicken, rat, mouse, human MRs. Maximum likelihood (ML) analysis was conducted using the JTT+G model. Statistical confidence for each branch in the tree was evaluated by the bootstrap method [41] with 1000 replications. We used the MEGA5 program [42] for these analyses.

### Construction of plasmid vectors

Full-coding regions of mineralocorticoid receptors were amplified by PCR with KOD DNA polymerase. PCR products were gel-purified and ligated into pcDNA3.1 vector. Mouse mammary tumor virus-long terminal repeat (MMTV-LTR) was amplified from pMSG vector by PCR, and inserted into pGL3-basic vector containing the *Photinus pyralis* lucifease gene. All constructs were verified by sequencing.

### Transactivation Assay

Human embryonic kidney 293 (HEK293) cells were used in the reporter gene assay. Transfection and reporter assays were carried out as described previously [33], except that we used PEI-max as transfection reagent [43]. All transfections were performed at least three times, employing triplicate sample points in each experiment. The values shown are mean ± SEM from three separate experiments, and dose-response data and EC50 were analyzed using GraphPad Prism.

### Statistical methods

Results are presented as mean ± SE (SEM) from three separate experiments. All multi-group comparisons were performed using one-way ANOVA followed by Bonferroni test. Dose-response data and EC50 were analyzed using GraphPad Prism. *P* < 0.05 was considered statistically significant.

## RESULTS

### Isolation of mineralocorticoid receptors from sturgeon and gar

We cloned sturgeon MR cDNA containing an open reading frame encoding 953 amino acids (GenBank accession LC149818)], and gar MR cDNA containing an open reading frame encoding 987 amino acids (GenBank accession LC149819). Sturgeon and gar MR sequences can be divided into four domains (Figure 3). The overall amino acid identity between these two MRs was 72%, with particularly high sequence identities for the DBD (100%) and LBD (89%) (Figure 3). Comparison of sturgeon MR with five other species (human, chicken, alligator, *Xenopus*, and zebrafish) revealed that sturgeon MR had identities of 44-36% in A/B domains,100-95% in DBDs, 67-47% in D domains, and 90-74% in LBDs (Figure 3).

**Figure 3.**
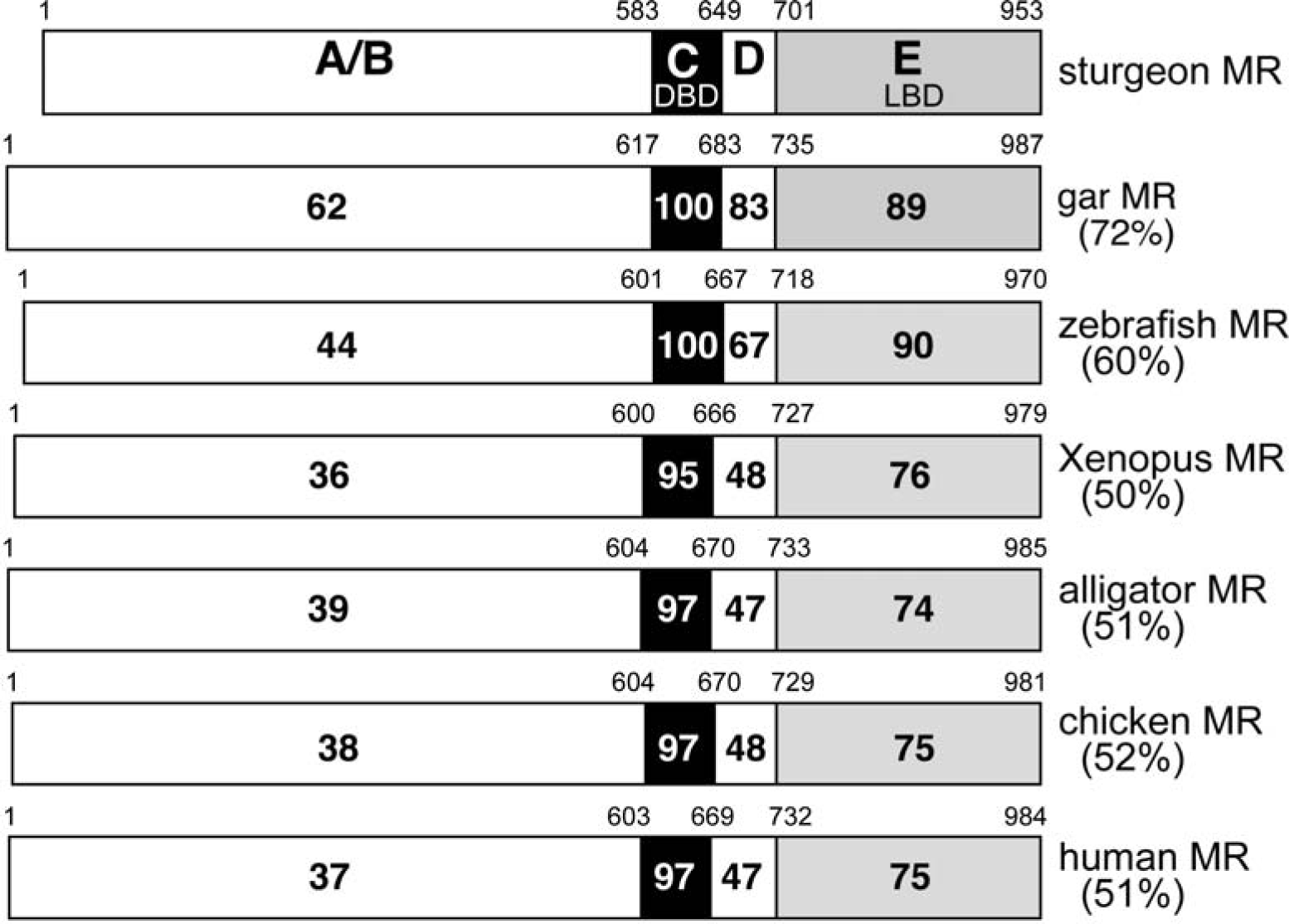
Comparisons of functional domains in sturgeon, gar, zebrafish, *X. laevis*, alligator, chicken and human MRs. Comparison of the domains in sturgeon MR gar, zebrafish, *X. laevis*, alligator, and human MR MR. The functional A/B domain, C domain, D domain and E domain are schematically represented with the numbers of amino acid residues at each domain boundary indicated. The percentage of amino acid identity between domains is depicted. GenBank accession numbers are: LC149818 for sturgeon MR; LC149819 for gar MR; NM_001100403 for zebrafish MR; NM_001090605 for *Xenopus* MR; AB701406 for alligator MR; and NM_000901 for human MR.

### Phylogenetic analysis of ancient fish corticoid receptors

To investigate the evolutionary position of gar and sturgeon MR in relationship to other fish MRs and tetrapods, we collected MR sequences from several teleosts, skates and elephant shark and selected terrestrial vertebrates. Consistent with the evolution of Acipenseriformes and Holostei, phylogenetic analysis places sturgeon and gar MRs close to the base of ray-finned fish (Figure 2).

### Strong response to 3-keto-steroids by sturgeon and gar mineralocorticoid receptors

We examined steroid-inducible transcriptional activation of gar and sturgeon MRs using the MMTV-driven reporter construct [33, 44]. For comparison, we also examined transcriptional activation of human MR and zebrafish MR. At 1 nM, Aldo, B, S, DOC, F and Prog were strong inducers of luciferase activation by gar MR and sturgeon MR and by zebrafish MR, with the exception of Prog which had a lower signal. These MRs show little stimulation by 1 nM DHT and E2 (Figure 4). At 1 nM, Aldo, B, DOC were strong transcriptional activators of human MR, which was activated to a lesser extent by S, and weakly activated by Prog (Figure 4).

**Figure 4.**
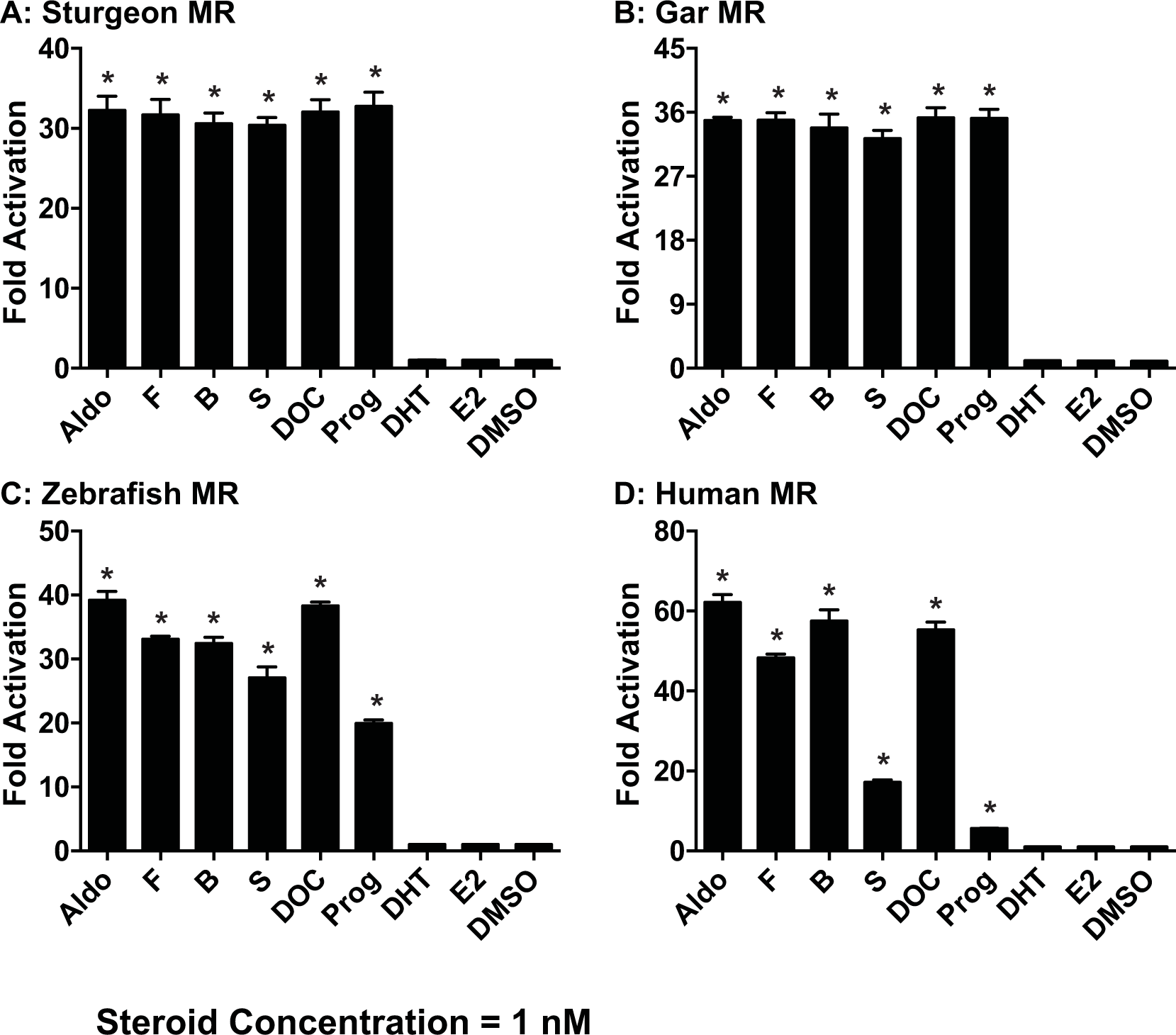
Ligand-specificities of fish and human MRs. Full-length sturgeon MR (A), gar MR (B), zebrafish (C), and human MR (D) were expressed in HEK293 cells with an MMTV-luciferase reporter. Cells were treated with 10^−8^ M Aldo, F, B, S, DOC, Prog, 5α-dihydrotestosterone (DHT), 17β-estradiol (E2) or vehicle alone (DMSO). Results are expressed as means ± SEM, n=3. Y-axis indicates fold-activation compared to the activity of control vector with vehicle (DMSO) alone as 1.

The agonist activity of Prog for sturgeon, gar and zebrafish MRs and for trout MR [16] and the evidence that spiron is an agonist for trout [16] and zebrafish MRs [19] stimulated us to examine transcriptional activation of sturgeon and gar MRs by spiron, an agonist for human MR. We also investigated transcriptional of sturgeon, gar and zebrafish MRs by 19norP, which, like Prog and spiron, is an agonist for human Ser810Leu-MR [27]. As shown in Figure 5, spiron and 19norP are agonists for sturgeon, gar and zebrafish MRs.

**Figure 5.**
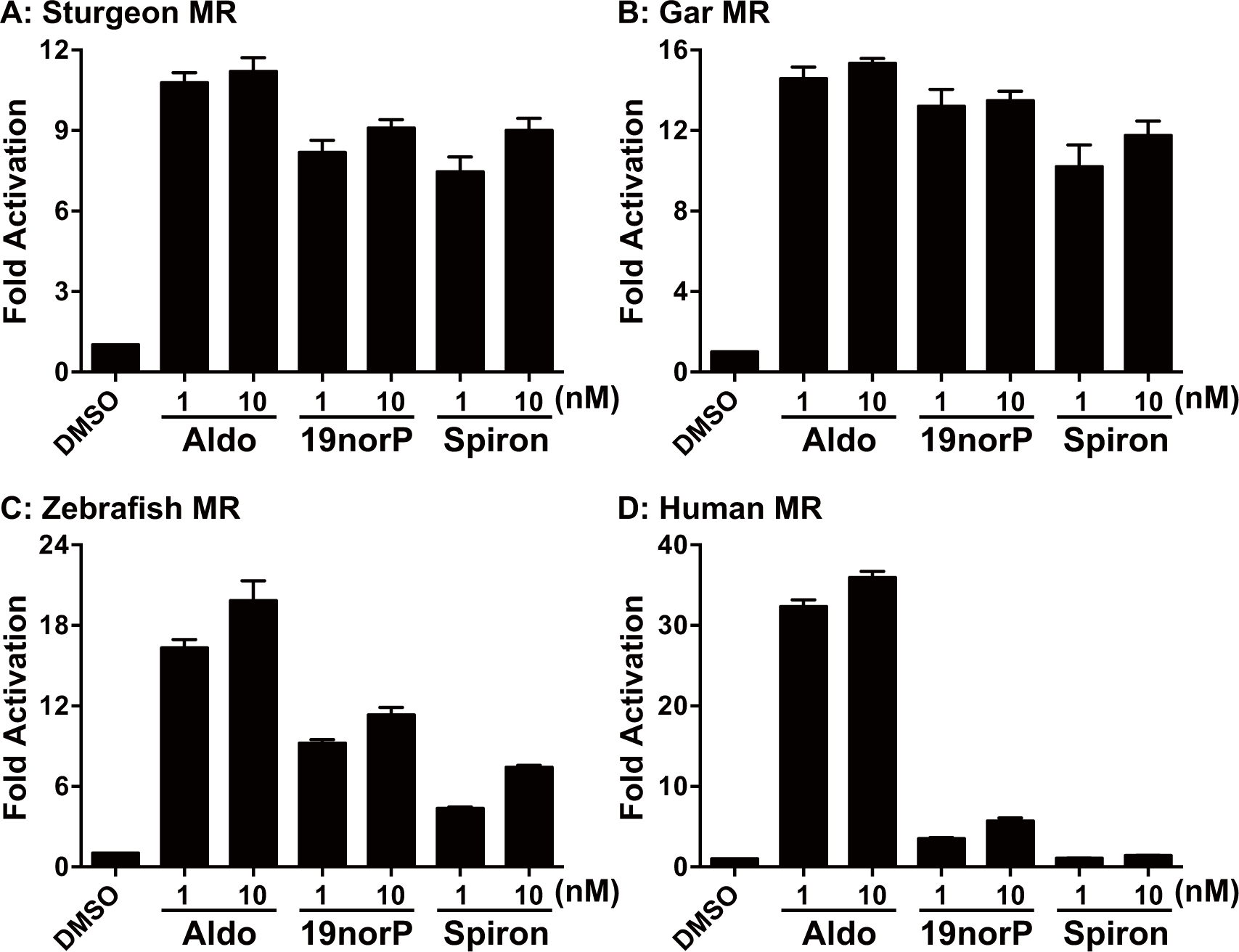
Transcriptional activation by 19nor-progesterone and spironolactone of sturgeon, gar, zebrafish and human MRs. Full-length sturgeon MR (A), gar MR (B), zebrafish MR (C), and human MR (D) were expressed in HEK293 cells with an MMTV-luciferase reporter. Cells were treated with 1 nM or 10 nM Aldo, 19norP or spiron or vehicle alone (DMSO). Results are expressed as means ± SEM, N=3 and represent fold-activation compared to the control vector with vehicle.

We examined concentration-dependent activation of gar, sturgeon, zebrafish, and human MRs by Aldo, F, B, DOC, S and Prog (Figure 6, Table 1). Both gar and sturgeon MRs had similar low EC50s, which varied from 7.7 pM to 150 pM for these steroids. For each steroid, the EC50s for gar MR were a little lower than for sturgeon MR.

**Figure 6.**
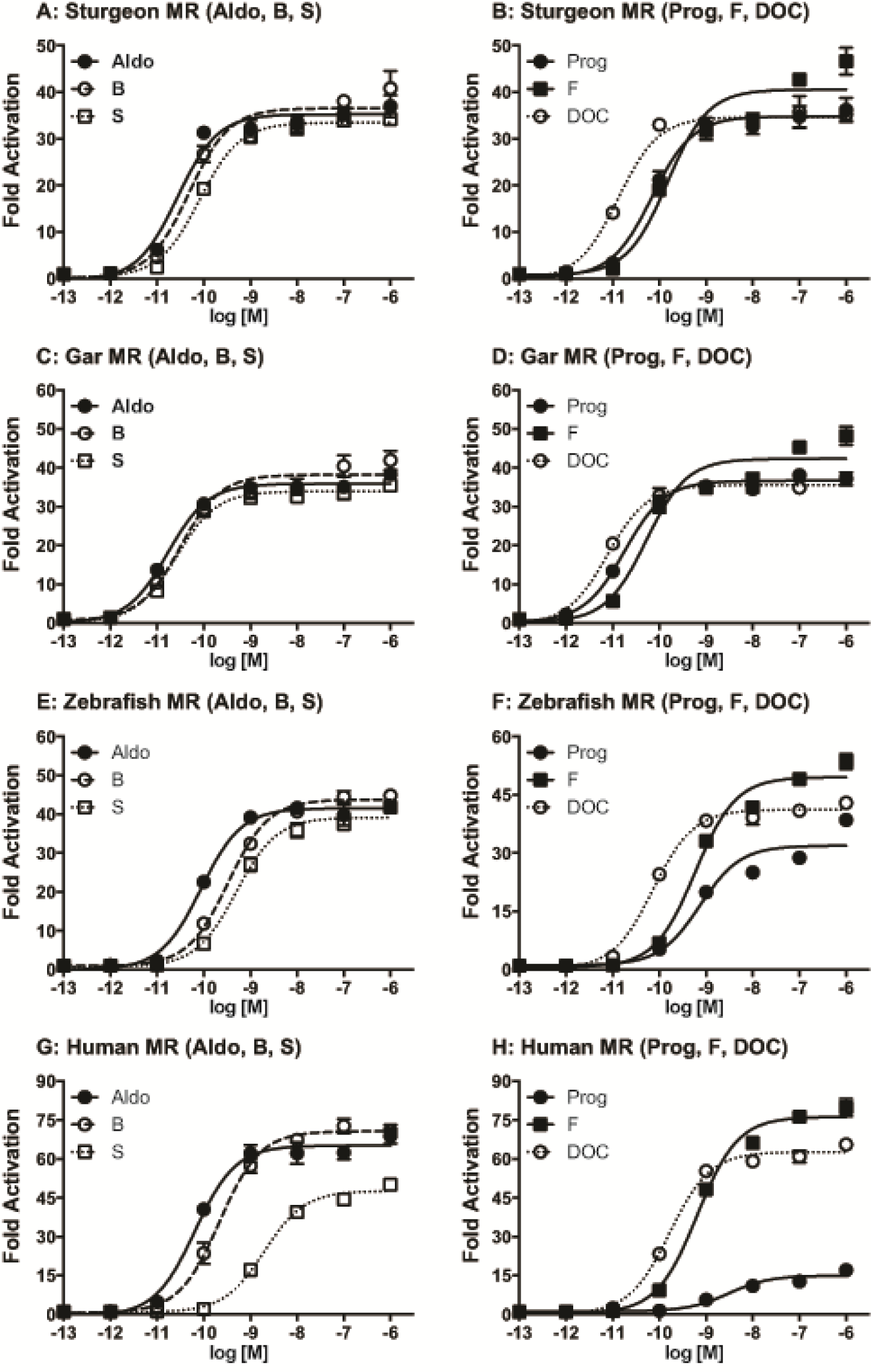
Concentration-dependent transcriptional activities of fish and human MRs. Concentration-response profiles of full-length sturgeon MR (A and B), gar MR (C and D), zebrafish MR (E and F), and human MR (G and H) for various steroids. HEK293 cells were transiently transfected with the MMTV-containing vector together with an MR expression vector. Cells were incubated with increasing concentrations of Aldo, B, and S (A, C, E, and G) or Prog, F, and DOC (B, D, F, and H) (10^−13^ to 10^−6^M). Data are expressed as a ration of steroid to vehicle (DMSO). Each column represents the mean of triplicate determinations, and vertical bars represent the mean ± SEM.

**Table 1.** EC50 activities for 3-keto-steroid transcriptional activation of sturgeon, gar, zebrafish and human MRs.

|  | Aldo | DOC | B | S | F | Prog |
| --- | --- | --- | --- | --- | --- | --- |
| Sturgeon MR | $2.7 \times 10^{-11}$ | $1.3 \times 10^{-11}$ | $4.8 \times 10^{-11}$ | $8.2 \times 10^{-11}$ | $1.5 \times 10^{-10}$ | $7.0 \times 10^{-11}$ |
| Gar MR | $1.7 \times 10^{-11}$ | $7.7 \times 10^{-12}$ | $3.1 \times 10^{-11}$ | $2.6 \times 10^{-11}$ | $5.3 \times 10^{-11}$ | $1.8 \times 10^{-11}$ |
| Zebrafish MR | $8.8 \times 10^{-11}$ | $7.4 \times 10^{-11}$ | $3.3 \times 10^{-10}$ | $5.0 \times 10^{-10}$ | $5.9 \times 10^{-10}$ | $7.4 \times 10^{-10}$ |
| Human MR | $6.5 \times 10^{-11}$ | $1.7 \times 10^{-10}$ | $2.2 \times 10^{-10}$ | $2.0 \times 10^{-9}$ | $6.5 \times 10^{-10}$ | - |

In comparison, EC50s of Aldo, B and F were similar for zebrafish and human MR and a little higher than their EC50s for sturgeon and gar MR. EC50s of DOC, S and Prog for zebrafish MR were higher than their EC50s for sturgeon and gar MR, but lower than the EC50s for human MR. Prog had a lower, but still significant, maximal activation for zebrafish MR while 100 nM Prog had little activation of human MR. Overall all corticosteroids and Prog had EC50s that would be consistent with a physiological role in transcription of the MR in sturgeon, gar and zebrafish (Table 1, Figure 6).

In human MR, Ser-810 and Ala-773 are important in the low transcriptional activity of Prog. Prog, spiron and 19norP can activate human MR with selective mutations at either Ser-810 or Ala-773 [27, 37, 38]. For example, at 1 nM, prog, spiron and 19norP are agonist for a Ser810Leu mutant MR [27, 37, 38]. We extracted the sequence of helices 3-5, which contain Ser-810 and Ala-773, from sturgeon, gar and zebrafish MR (Figure 7) and other teleosts. All of these ray-finned fish contain a serine and alanine that aligns with Ser-810 and Ala-773 in human MR.

**Figure 7.**
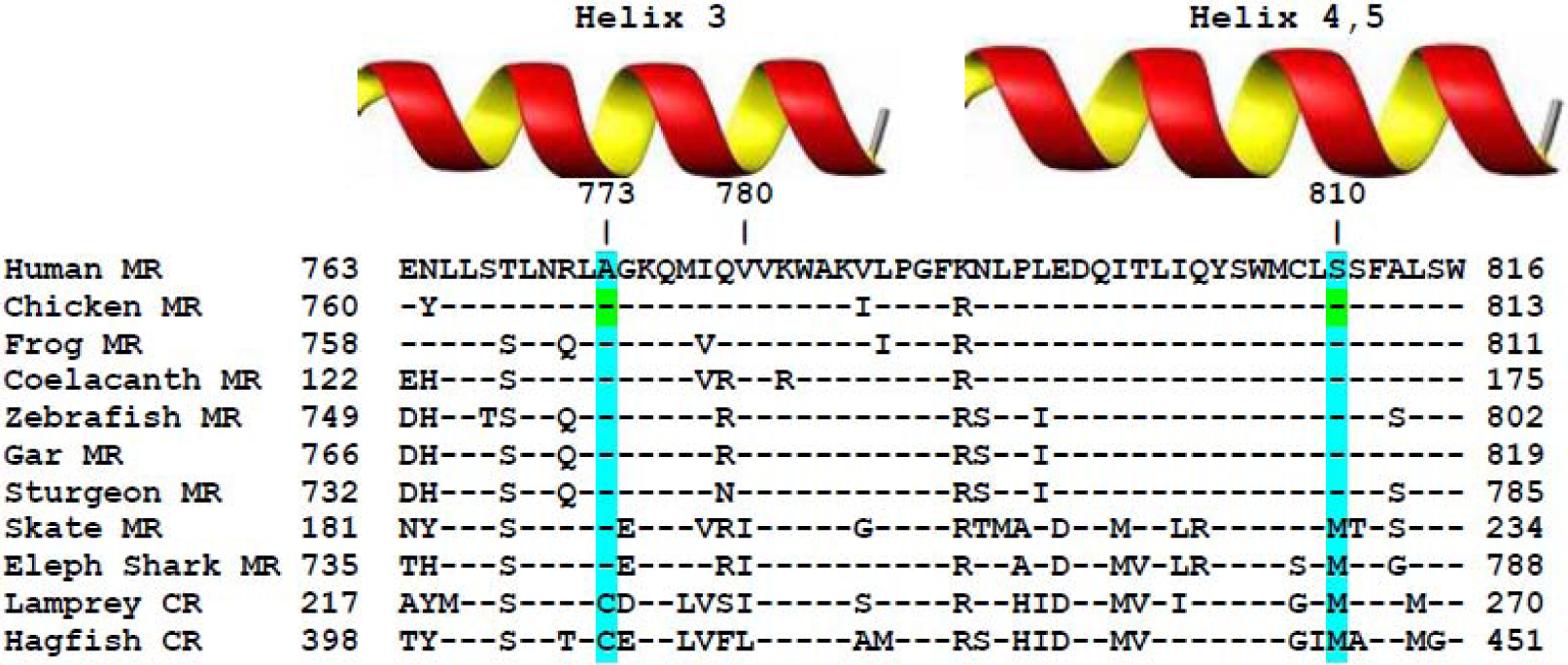
Alignment of vertebrate MRs to Serine-810 and Alanine-773 in helices 3-5 in human MR. Human Ser-810 and Ala-773 are conserved in ray-finned fish MRs. Skate MR, elephant shark MR, lamprey CR and hagfish CR contain a methionine corresponding to human Ser-810. Lamprey CR and hagfish CR contain a cysteine corresponding to Ala-773 in human MR. Amino acids that are identical to amino acids in human MR are denoted by (-).

## DISCUSSION

Several corticosteroids are physiological activators of the MR in cartilaginous fishes, ray-finned fishes and tetrapods. Interestingly, Aldo, the mineralocorticoid for terrestrial vertebrates, first appears in lungfish [45]. Nevertheless, Aldo is a potent activator of the lamprey CR [6], which is ancestral to the MR [5-7, 11]. Interestingly, F, DOC, B and S and Prog also are transcriptional activators of the CR in lamprey and hagfish [6], with only S, thus far, found to have mineralocorticoid activity in lamprey [12, 13]. In skate, which has separate MR and GR genes, Aldo, F, DOC and B are strong transcriptional activators of the MR [14]. F, DOC, B, S and Prog are found in teleosts [26] and F and DOC have been proposed to be transcriptional activators of teleost MRs [4, 15-19, 21, 22, 24, 25].

Absent, until now, was information about the response to corticosteroids of MRs in sturgeon and gar, two basal fish that fill in the gap between elasmobranchs and teleosts (Figure 2). Here we report that sturgeon MR and gar MR have EC50s below 1 nM for Aldo, F, DOC, B, S and Prog. Interestingly, we find that zebrafish MR also has a similar strong response to these corticosteroids and Prog. Moreover, spiron and 19norP are agonists for sturgeon, gar and zebrafish MRs. This low selectivity for 3-keto-steroids (Figure 1) that can activate these fish MRs resembles the response to these steroids by lamprey and hagfish CR [6] and skate MR [14]. Thus, this strong response of the MR to a broad panel of 3-keto-steroids was conserved after the third whole-genome duplication at the base of the teleosts [30-32, 46].

In contrast, human MR is more selective for 3-keto-steroids with higher EC50s for S and Prog. Our data showing weak activation by Prog and 19norP of human MR is in agreement with other studies [35-37]. The strong response to Prog, spiron and 19norP of ray-finned fish MR is interesting in the light of the report by Geller et al. [27] that human MR with a Ser810Leu mutation was activated by 1 nM Prog, 19norP and spiron. Mutagenesis studies and structural analyses of the MR-Leu810 mutant led to the hypothesis that Leu-810 on α-helix 5 has stabilizing van der Waals contacts with Ala-773 on α-helix 3 [27, 37, 38] to explain the strong transcriptional activation by Prog, 19norP and spiron. This serine and alanine are conserved in and sturgeon and gar MRs, as well as in zebrafish MR (Figure 7) [4] and other teleost MRs indicating that other mechanism(s) can lead to a strong response of sturgeon, gar and zebrafish MR to Prog, 19norP and spiron. Activation by Prog of zebrafish MR is of concern because zebrafish is an established model system for studying gene regulation in teleosts, as well as providing insights into human physiology [47]. Prog activation of zebrafish MR may confound data that focuses on activation of the PR. Prog may also be an agonist for the MR in medaka and other teleosts that have a serine and alanine that correspond to Ser-810 and Ala-773 in human MR.

### Mechanisms for regulation of steroid activation of ray-finned fish MR

The strong response of zebrafish MR, as well as sturgeon and gar MRs, to five corticosteroids, Prog, 19norP and spiron requires one or more mechanisms to provide steroid-specific regulation of transcriptional activation of these ray-finned fish MRs. At this time, such mechanisms in gar, sturgeon and zebrafish MRs or other ray fined fish MRs are poorly understood. Clues for possible mechanisms may be found from insights into regulation of mammalian MRs [4, 11, 48-53]. One possibility is an important mechanism in epithelial cells for regulating access of F and B to mammalian MR by tissue specific expression of 11β-HSD2, which selectively converts F and B, respectively, to cortisone (E) and 11-dehydrocortisone (A), two inactive steroids. Aldo is inert to 11β-HSD2, allowing Aldo to occupy the MR in epithelial cells in which 11β-HSD2 inactivates F and B [11, 51, 52]. 11β-HSD2 is found in ray fined fish [54, 55], including sturgeon (Accession: ALE30175) and gar (Accession: XP_006641583). Expression of 11β-HSD2 in MR-containing tissues provides a mechanism to exclude F and B from the MR. DOC, S and Prog, which have low EC50s in gar, sturgeon and zebrafish, lack an 11β-hydroxyl group and are inert to 11β-HSD2.

Other regulatory mechanisms of the response of the MR to 3-keto-steroids include tissue-selective synthesis of 3-keto-steroids [5, 56-58], selective sequestration of 3-keto-steroids to plasma proteins [48, 53, 59], steroid-specific conformational changes that regulate MR binding of co-activators [60-64], effects of inter-domain interactions between the NTD and the LBD [19, 50, 64, 65] and post-translational modifications, such as phosphorylation, and SUMOylation [49, 66, 67].

## Authors Contributions

A.S., K.O., R.S., and S.A. carried out the research. M.E.B. and Y.K. conceived and designed the experiments and wrote the paper. All authors gave final approval for publication. We have no competing interests.

## Acknowledgments

We thank colleagues in our laboratories. We also thank Drs. Arlette Hernandez-Franyutti and Mari Carmen for providing gar tissues.

